# Intracellular delivery of mRNA to human primary T cells with microfluidic vortex shedding

**DOI:** 10.1101/343426

**Authors:** Justin A. Jarrell, Amy A. Twite, Katherine H. W. J. Lau, Moein N. Kashani, Adrian A. Lievano, Julyana Acevedo, Craig Priest, Jorge Nieva, David Gottlieb, Ryan S. Pawell

## Abstract

Intracellular delivery of functional macromolecules, such as DNA and RNA, across the cell membrane and into the cytosol, is a critical process in both biology and medicine. Herein, we develop and use microfluidic chips containing post arrays to induce *microfluidic vortex shedding*, or *μVS*, for cell membrane poration that permits delivery of mRNA into primary human T lymphocytes. We demonstrate transfection with *μVS* by delivery of a 996-nucleotide mRNA construct encoding enhanced green fluorescent protein (EGFP) and assessed transfection efficiencies by quantifying levels of EGFP protein expression. We achieved high transfection efficiency (63.6 ± 3.44% EGFP+ viable cells) with high cell viability (77.3 ± 0.58%) and recovery (88.7 ± 3.21%) in CD3+ T cells 19 hrs after *μVS* processing. Importantly, we show that processing cells via *μVS* does not negatively affect cell growth rates or alter cell states. We also demonstrate processing speeds of greater than 2.0 × 10^6^ cells s^−1^ at volumes ranging from 0.1 to 1.5 milliliters. Altogether, these results highlight the use of *μVS* as a rapid and gentle delivery method with promising potential to engineer primary human cells for research and clinical applications.

## Introduction

Biomicrofluidics are used to isolate^1^, enrich^1^, modify^2,3^, culture^4^ and qualify cells^5^, lending to the development and manufacturing of gene-modified cell therapy (GMCT) where these processes are vital. GMCTs based on chimeric antigen receptor-expressing T-cells (CAR-T) can provide substantial improvement in patient outcomes, including complete remission of disease for hematologic malignancies^6^. CAR-T cells targeting CD19, for example, have demonstrated 83% clinical remission in patients with advanced acute lymphoblastic leukemia who were unresponsive to prior therapies^7^. These unprecedented results exemplified in multiple clinical trials have made CD19-targeting GMCT the first to gain approval by the FDA^7^.

The current standard for manufacturing GMCTs involves using viral-based gene transfer which is costly, time consuming, and can have variable results^8–10^. In addition, viral transduction for CAR-T therapies requires extensive safety and release testing for clinical development and post-treatment follow-up^9^. Unlike viral-based methods, electroporation can be used to deliver a broader range of bioactive constructs into a variety of cell types, while bypassing the extensive safety and regulatory requirements for GMCT manufacturing using viruses^8,9^. However, the significant reductions in cell numbers and viabilities, accompanied by changes in gene expression profiles that negatively impact cell function, make physical transfection methods like electroporation less than ideal for GMCT applications^2,3,9,11–13^. Therefore, the ideal intracellular delivery method to generate GMCTs would permit transfection of various constructs to multiple cell types while having minimal effects on cell viability and cell recovery, and minimal perturbation to normal and/or desired (i.e. therapeutic) cell functions^2,3^.

In general, microfluidic methods have improved macromolecule delivery into cells by scaling microfluidic channel geometries with cell dimensions. Intracellular delivery methods utilizing microfluidics include electroporation^14–16^, microinjection^17^, cell constriction or squeezing^18–23^, fluid shear^24,25^ and electrosonic jet ejection^26,27^. These methods offer appealing alternatives to conventional transfection systems, however, their production output (i.e. number of engineered cells) is limited by throughput, processing speeds, and clogging as a result of cell shearing, cell lysis, and debris formation^2,3^. Thus, it remains unclear as to how well these methods may scale for clinical-level production of GMCTs that often require greater than 10^7^-10^8^ cells per infusion^28,29^.

There are several practical metrics when considering microfluidic intracellular delivery for GMCTs including cell viability, cell recovery, delivery or expression efficiency, sample throughput, and cell states and functions. Importantly, GMCTs require large numbers of viable, gene-modified cells to enhance clinical response rates and prevent adverse events in patients^28,29^. For instance, infusion of genetically-modified, non-viable cells have been shown to promote toxicities *in vivo*, requiring additional safety and regulatory considerations if present in high numbers prior to treatment^30–32^. Additionally, in terms of processing, low throughput transfection methods, such as single cell micro-needle injection, would require 10^8^ seconds, or three years, to deliver a single autologous GMCT without expansion^,17,28,29^. To overcome these limitations, next generation transfection methods need to preserve high cell viabilities and desired cell states prior to infusion, while implementing strategies to increase processing throughput (i.e. sample size, number and processing speed), diminish production times, and minimize processing steps. As it stands, the current state of microfluidic and mechanical intracellular delivery methods described above fail to meet the needs of GMCT development and clinical-level manufacturing.

Microfluidic vortex shedding, or *µVS*, is based on a well-known macroscale phenomenon^33^ known to occur in dense arrays of microfluidic posts^34^. Current transfection methods that use mechanical microfluidics as the method of delivery either do not use post arrays (i.e. microfluidic jet ejections) ^26,27^ or use arrays with spacing smaller than a cell’s diameter (i.e. squeezing)^18–23^ that dramatically reduce processing rates. Interestingly, the hydrodynamic conditions that facilitate *µVS* in a microfluidic post array with spacing greater than a cell’s diameter suggests that our device can efficiently deliver material into cells while addressing the limitations of physical transfection methods. Therefore, we sought to implement *µVS* in the construction of a device to deliver mRNA into cells.

Here, we describe the development and evaluation of our microfluidic device for hydrodynamic, intracellular delivery of mRNA into human T cells using *μVS*. As a proof-of-concept of our technology, we delivered a 996-nucleotide mRNA construct expressing enhanced green fluorescent protein (EGFP) into primary human T lymphocytes and demonstrated high levels of cell viability, cell recovery, and EGFP expression with minimal effects to cell growth and activation profiles after delivery. We demonstrate that *μVS* does not adversely affect T cell growth, results in high transfection efficiencies, high cell viability and even expression profiles among CD4+ and CD8+ T cells after transfection at processing rates exceeding 2 × 10^6^ cells s^−1^.

## Results

### Empirical Verification of Microfluidic Vortex Shedding (*μVS*)

Vortex shedding has been previously shown to occur in microfluidic post arrays^34^. Therefore, we iterated through various engineering features, including post sizes and diameters, post shapes, post-array spacing, number of rows within the post-array region and fluid pressure to generate a prototype chip design for intracellular delivery of mRNA to human T cells (data not shown). Our results indicated that six rows of circular posts of 40 μm diameter with spacing between each post approximately twice the diameter of a human T cells (~9.0μm, 8 to 12μm) and an applied gauge pressure of 120 PSI was ideal for delivery of mRNA (Figure 1c-d) to T cells from a single donor when using compressed nitrogen. We manufactured chips based on these parameters and used them for transfection experiments with activated human T cells (Figure 1b).

**Figure 1.**
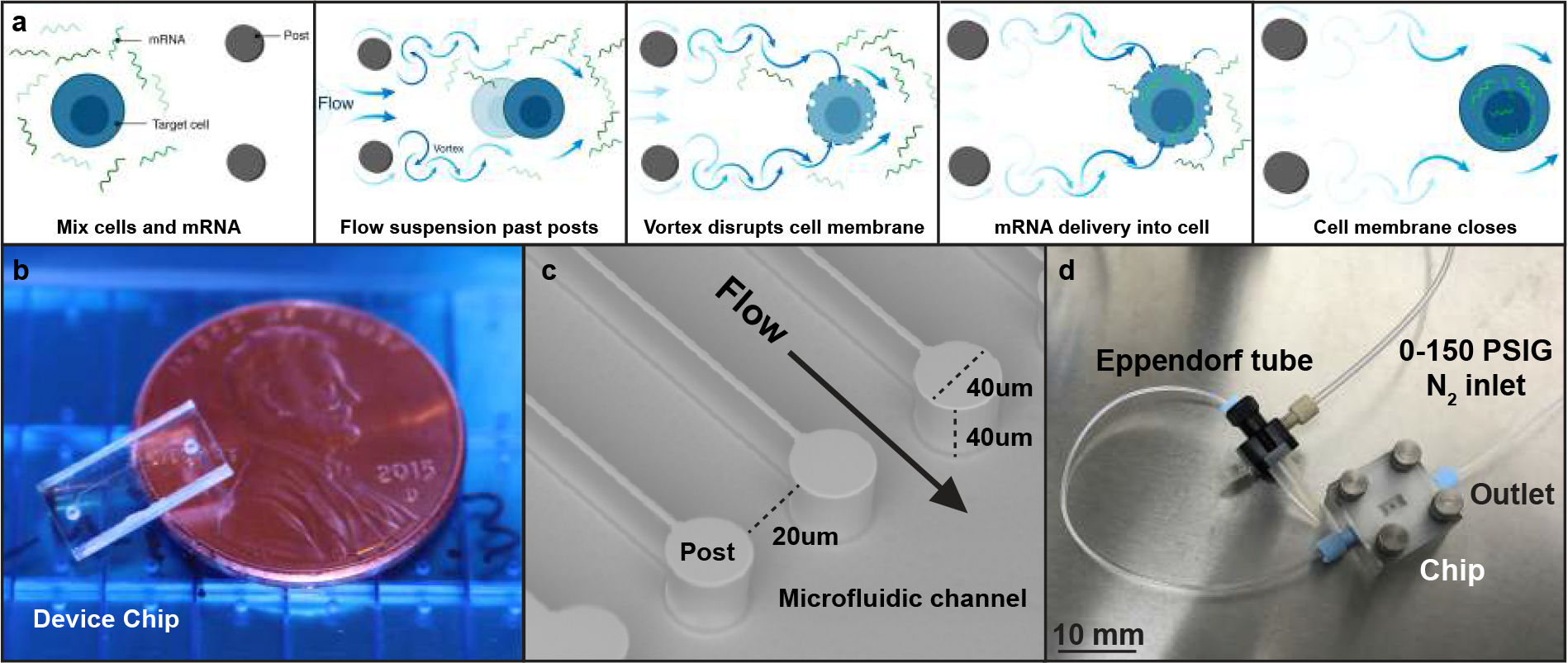
Device design, dimensions, and processing overview. (**a**) Schematic of microfluidic vortex shedding (*μVS*)- based delivery system used to transfect mRNA into human T lymphocytes. Briefly, cells and mRNA were mixed in suspension, suspension was flowed past ion etched posts creating vortices that disrupt cell membranes to permit poration and allow mRNA to diffuse into cells prior to membrane repair. (**b, c**) Image of microfluidic chip and channel highlighting flow direction and distance between posts that permit cells to pass through with minimal mechanical deformation (i.e. cell squeezing). The distance between posts is adjustable (10 – 40 microns) with a 20 micron distance was selected for this study based on cell type and size. (**d**) Image of hardware unit used to pneumatically push cell and mRNA suspension through chip. Schematic (a) is for illustrative purposes only and not drawn to scale.

*μVS* leverages naturally-occurring fluid dynamics to permeabilize cell membranes that may also lyse cells^2,3^. Therefore, it was also necessary to evaluate if build-up caused by cell debris resulted in constriction-based cell poration, which may be the cause of any transfection not accounted for by *μVS*. If the primary method of transfection was constriction, we reasoned that samples collected at the end of an experiment would show significantly higher transfection efficiencies compared to samples collected at the beginning with less debris build up in the fluidic device and/or chip. To assess this, we processed human T cells through our device in the presence of EGFP-encoding mRNA and quantified the level of EGFP protein expression in cells collected at various time points. Samples collected at the start of the experiment had EGFP expression levels comparable to samples collected at the end, with no significant differences observed across all time points (n.s. for all comparisons, Supplemental Figure 1). Our observation of consistent EGFP expression across all time points during processing suggested that constriction-based deformation was not contributing to the transfection observed in our system. We conclude that any build up accumulated while processing samples through our device has a negligible impact on transfection efficiency, with cell poration and mRNA delivery attributed primarily to *μVS*.

### Hydrodynamic Characterization

*μVS* is a hydrodynamic phenomenon shown to occur in microfluidic post arrays at an object Reynold’s number (Re_o_) > 40^34^. To determine if the hydrodynamic conditions required to induce and sustain vortex shedding are achieved in our flow cells, we observed and characterized flow dynamics using non-dimensional analysis and computational fluid dynamic simulations. Since our processing media was largely composed of water, we characterized hydrodynamic conditions using the kinematic viscosity of water as 20 °C (1.004 × 10^−6^ m^2^ s^−1^). Our characterization specified an Re_o_ = 146 around posts at typical experimental conditions, indicating that the hydrodynamic conditions within the post-array regions of our flow cells were in the range of *μVS*. A summary of our non-dimensional characterization is shown on Table 1 with calculations outlined in the *SI*.

**Table 1.** Non-dimensional hydrodynamic characterization of μVS

| Feature | Units | Value |
| --- | --- | --- |
| Flow cell Reynold's number ($Re_{fc}$) | — | 271 |
| Channel Reynold's number ($Re_c$) | — | 180 |
| Gap Reynold's number ($Re_g$) | — | 291 |
| Object (Post) Reynold's number ($Re_o$) | — | 146 |
| Strouhal number ($St$) | — | 0.18 |
| Frequency ($f$) | kHz | 14.8 |

Computational fluid dynamic simulations were used to advance our non-dimensional analysis of hydrodynamic flow conditions and determine the velocity magnitude and direction of induced vortex shedding and Q-criterion for vortex development in our flow cell. Flow conditions in the simulation were selected to represent experimental conditions. Simulations of the contour of velocity magnitude and vector showed the formation of vortices between posts of the post-array, with flow separating from the post surfaces at high velocities (Figure 2a-b). Significant decreases in flow velocity were observed in the fluid exiting the post-array region into exit channels coupled with the resolution of vortices (Figure 2b). To further assess vortex formation, Q-criterion magnitudes were analyzed at various regions in our flow simulations, with the highest values detected in regions immediately after each post-array row where vortex development occurs (Figure 2c). Conversely, near zero Q-criterion values were observed within the inlet channels in the absence of vortex development, with comparable values observed in the exit channels as vortices resolved (Figure 2c). Similarly, flow conditions in a blank chip without post-arrays showed zero Q-criterion along the entire channel (Supplemental Figure 2). Altogether, these results support that the hydrodynamic conditions observed in our flow cell induce *μVS.*

**Figure 2.**
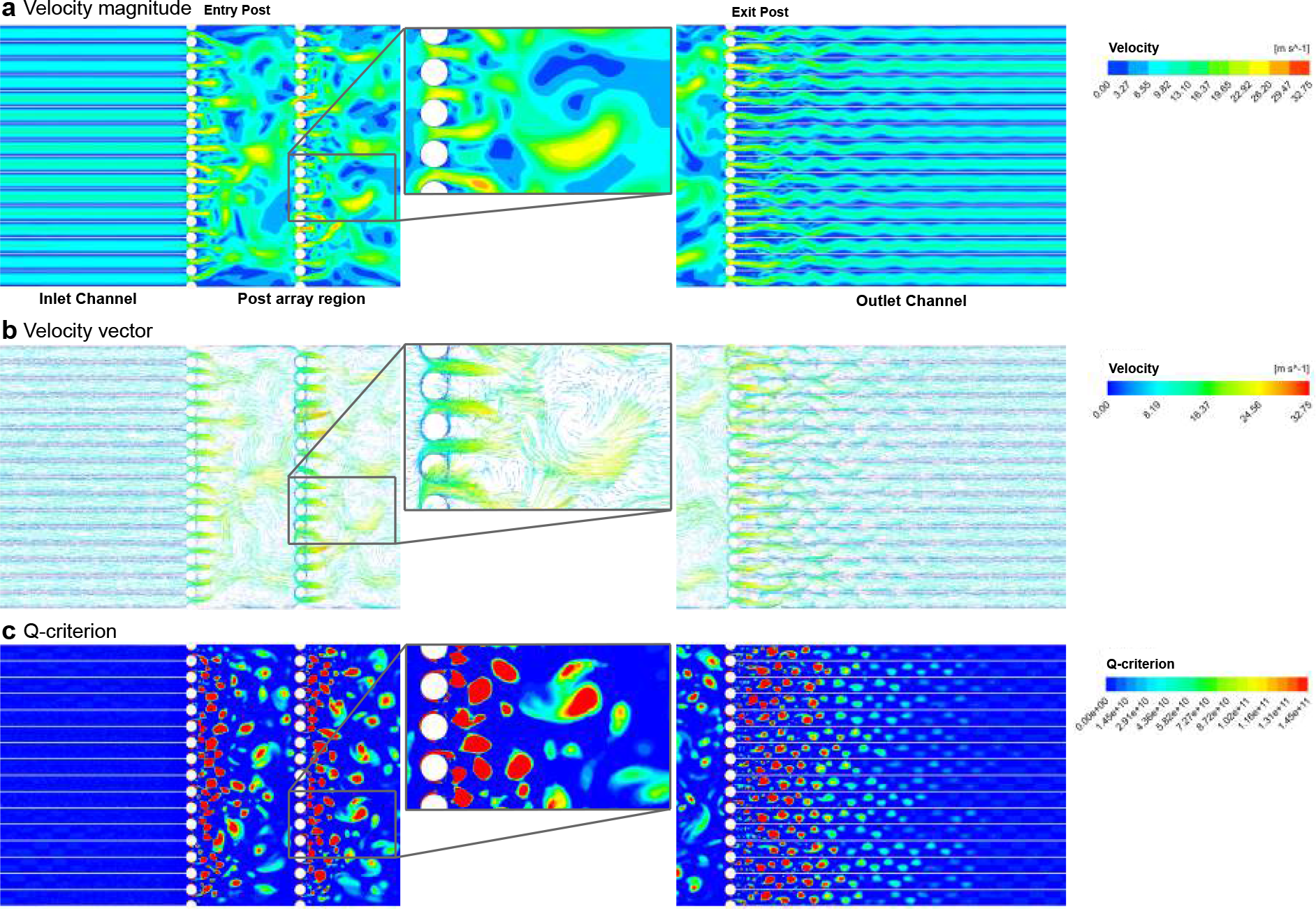
Computational fluid dynamic simulation of on-chip microfluidic vortex shedding. Simulation of the flow dynamics in a 960μm width × 6.95mm length chip containing 40μm diameter posts depicting (**a**) velocity magnitude (m s^2^), (**b**) velocity vector (m s^2^), and (**c**) Q-criterion. Flow direction was simulated from left to right. Images show inlet and outlet channels and post-array region containing 6 rows of posts, with the 1^st^ (entry) and 6^th^ (exit) post rows highlighted. Vortices of various sizes are shown in enlarged insets.

### EGFP mRNA delivery into primary human T cells using *μVS*

Macroscale electroporation and viral transduction are effective methods for intracellular delivery into T cells^2,3,6–12,35–38^. For example, delivery of an EGFP-encoding plasmid in cultured peripheral mononuclear cells via electroporation resulted in >60% EGFP expression in CD3+ T cells post transfection^11^. Therefore, we sought to investigate if comparable transfection efficiencies (>60%) could be achieved in human T cells via *μVS*.

For our proof-of-concept study, we selected an approximately 1.0kB EGFP mRNA construct to transfect into primary human CD3+ T cells via *μVS*. We observed an increased level of EGFP expression at mRNA concentrations ranging from 10 to 160 μg mL^−1^ compared to cells processed through our device without EGFP mRNA (‘processing control’), with 160μg mL^−1^ EGFP mRNA without device processing (‘mRNA control’), or without mRNA or device processing (‘handling control’) (Figure 3a). EGFP expression levels ranging from 23.6% to 63.6% in live cells were observed approximately 19 hours post-transfection, with a statistically significant increase in expression detected in cells processed with 160 μg mL^−1^ EGFP mRNA (p <0.05, Figure 3a-b). Interestingly, the median EGFP fluorescence intensity observed in the live cell population was linearly correlated with mRNA concentration (R^2^ = 0.988, Figure 3c), indicating that the overall expression level of EGFP on each cell increased with higher concentrations of EGFP mRNA delivered via *μVS*.

**Figure 3.**
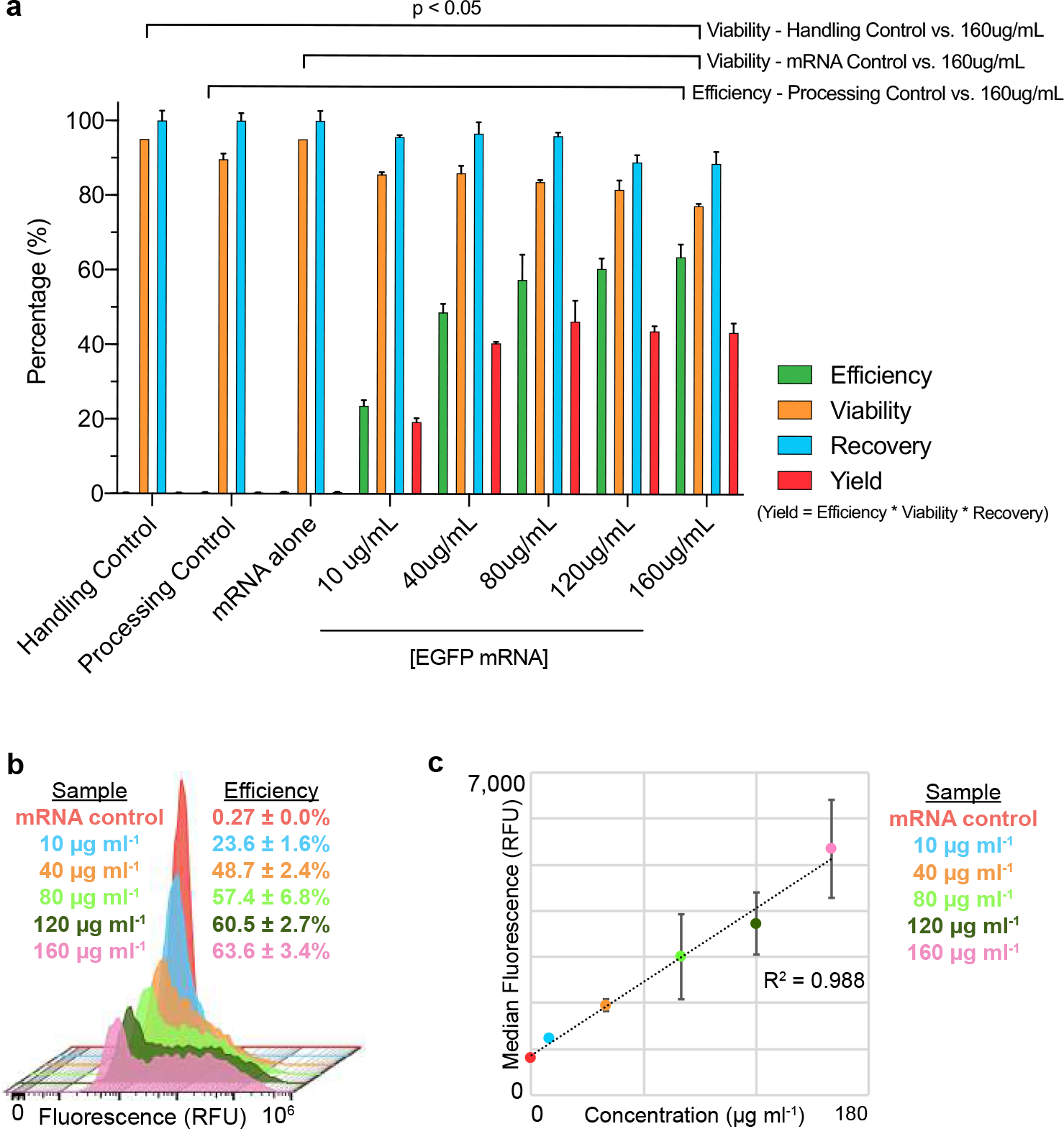
EGFP mRNA delivered to primary human T lymphocytes via *μVS*. (**a**) Levels of EGFP protein expression (Efficiency, green), cell viability (Viability, yellow), cell recovery (Recovery, blue), and total yield (Yield, red) with increasing concentration of 996 nt EGFP mRNA (10 to 160 μg mL^−1^) delivered via *μVS* compared to a handling control (no mRNA, no device processing), mRNA control (160 μg mL^−1^ EGFP mRNA without device processing), and *μVS* device processing control (no mRNA) at 19 hours post transfection (n = 3 per condition). (**b**) Histogram overlays of EGFP protein expression levels from live, single CD3+ T cells analyzed via flow cytometry as a function of mRNA concentration at 19 hours post transfection. (**c**) Scatter plot of the median EGFP fluorescent intensity (in relative fluorescence units, RFU) and mRNA concentration at 19 hours post transfection. R-squared value calculated and displayed on graph. Data represent the mean ± SD (a, b) or median ± SD (c) of triplicate values. P values were calculated between groups by one-way ANOVA and post-hoc Kruskal-Wallis multiple comparisons test (a).

Substantial reductions in cell viability and recovery are consistently observed in human T cells processed with rapid transfection methods like electroporation, with cell viabilities ranging from 15% to 40% for T cells immediately after processing^11,35,38^. Comparatively, we observed cell viabilities ranging from 86% to 77% when delivering 10 to 160 μg mL^−1^ of EGFP mRNA, respectively, compared to approximately 90% viability measured in controls (Figure 3a). A significant decrease in the percentage of viable T cells was observed at 160μg mL^−1^ mRNA, with no significant differences detected at concentrations from 10 to 120 μg mL^−1^ (p<0.05 for 160μg mL^−1^, Figure 3a). Similarly, no significant decreases in the percentage of cells recovered after *μVS* were observed for all concentrations of mRNA, ranging from 87% to 97% (n.s., Figure 3a). To take all measurements into consideration, we created a metric representing the product of the percentage of EGFP-expressing cells, viable cell, and recovered cells, and calculated the ‘yield’ for each experimental condition and controls. Comparison of the calculated values revealed a higher percent yield across all concentrations of mRNA delivered via *μVS* compared to controls (0.30% to 0.49%), with the highest yield (46.3%) observed at 80μg mL^−1^ mRNA (Figure 3a). Cumulatively, these results suggest that delivery of mRNA at various concentration via *μVS* varies the transfection efficiency, cell viability, and cell recovery, with optimal levels achieved when delivering ≥ 80μg mL^−1^ mRNA to previously activated human T cells.

Accompanying the substantial decrease in viability, cell death is considerable in electroporated cells and progresses days after transfection^11^. In comparison, intracellular delivery of mRNA via *μVS* showed only slight decreases in cell viability two days after transfection for all mRNA concentrations, ranging from 75% to 81% of viable cells compared to an average of 92% in controls (Figure 4a). Thereafter, *μVS*-transfected cells recovered rapidly with no significant differences in cell viabilities detected at day 3 and 1 week after transfection (n.s. for all comparisons with controls, Figure 4a, Supplemental Table 1). Furthermore, we did not observe any significant difference in the growth rates of transfected cells for all concentration of mRNA delivered to human T cells compared to controls (n.s. for all comparisons with control, Figure 4b). Interestingly, comparison of the processing and handling control conditions revealed near identical cell viabilities and growth rates across seven days, suggesting that decreases in these parameters are due to the presence of mRNA, oppose to *μVS* processing (Figure 4a-b). Quantifying the EGFP expression levels in transfected T cells revealed a peak transfection efficiency of 63.6% when delivering 160μg mL^−1^ of mRNA at 19 hours post-transfection, with levels decreasing to 39.9% of cell expressing EGFP by day 7 (Figure 4c). We found that the levels of EGFP expression were significantly higher across all mRNA concentrations and time points compared to handling and processing controls (p < 0.05 for all comparisons, Figure 4c, Supplemental Table 1). Altogether, these data indicate that *μVS*-based transfection is a quick (~3 s for 400 μL) and gentle method for intracellular delivery of mRNA into human T cells resulting in high transfection efficiency, cell viability and cell recovery.

**Figure 4.**
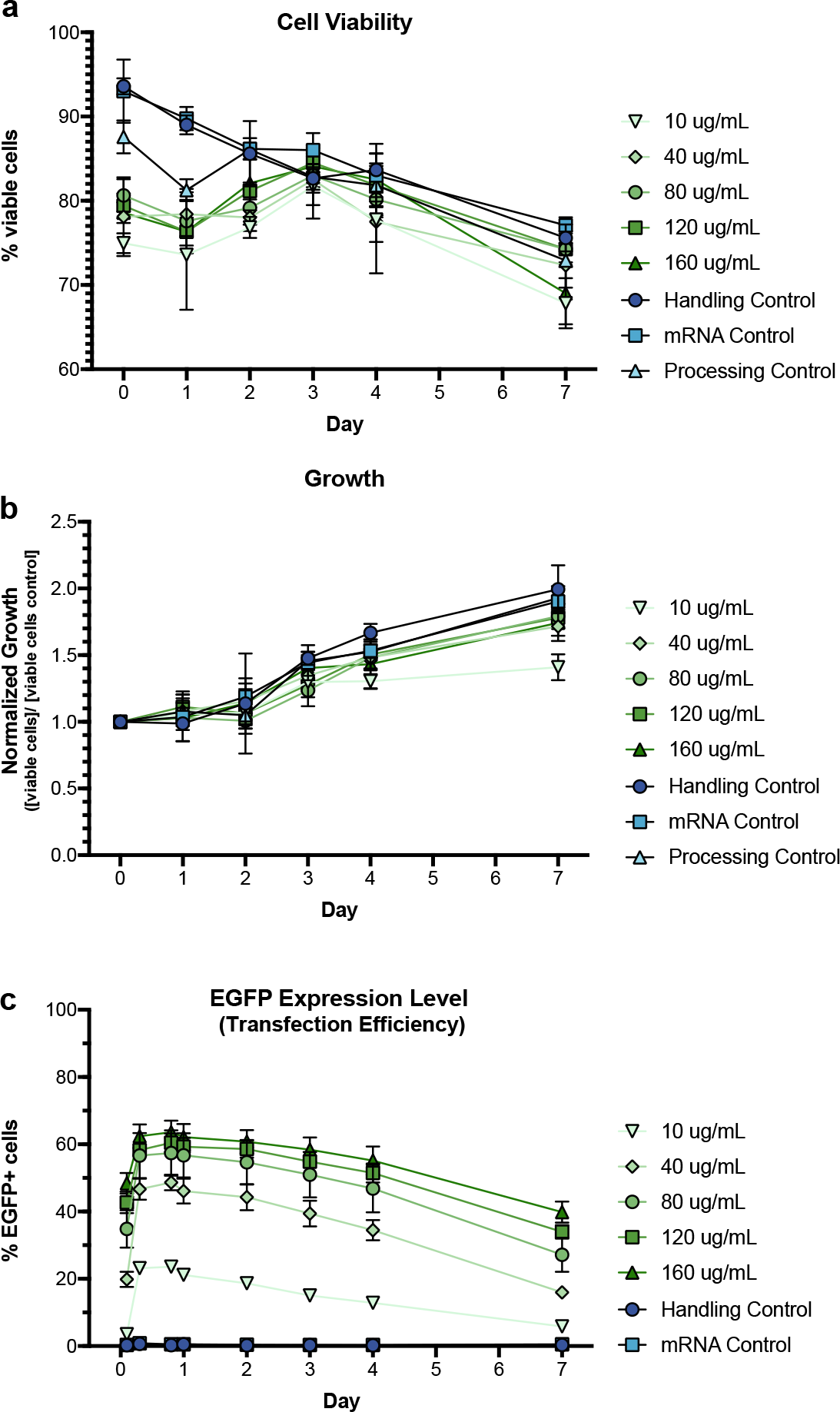
Cell viability, growth, and EGFP protein expression in primary human T lymphocytes transfected with EGFP mRNA via *μVS.* (**a**) Cell viability (% of viable cells over total cells), (**b**) normalized growth ([viable cells] per condition over [viable cells from handling control] per day), and (**c**) EGFP expression levels (% of EGPF+ cells over total cells) in human T cells transfected with increasing concentrations (10 to 160 μg mL^−1^) of EGFP mRNA via *μVS* compared to controls at various time points for 1-week post-transfection. Data represent means ± SD of triplicate values. P values were calculated between groups by two-way ANOVA and post-hoc Tukey’s multiple comparisons test (Supplemental Table 1).

### EGFP expression levels are equal among CD_4+_ and CD8+ T lymphocytes

Lentiviral- and electroporation-based delivery efficiencies can differ among T cell populations without a consistent trend between donors^11,36^. To quantify the expression level of EGFP in different T cell populations after *μVS*, we enriched human CD3+ T cells, transfected with three concentrations of EGFP mRNA (10, 80 and 160 μg mL^−1^ mRNA), and analyzed the CD4+ and CD8+ populations by flow cytometry. We observed a 4:1 distribution of CD4+ to CD8+ T cells of the total CD3+ population after *μVS* delivery (Figure 5). Interestingly, we detected near identical percentages of EGFP-expressing cells among CD4+ and CD8+ T cells of the total CD3+ population after *μVS* for all concentrations of mRNA post transfection (n = 3, Figure 5a-c). These expression levels ranged from 23.5% to 63.5% with the highest levels detected after delivery with 160 μg mL^−1^ EGFP mRNA, similar to the levels observed in our time course experiments (Figure 4c, 5c). Cumulatively, these results revealed an even level of EGFP expression in CD4+ and CD8+ human T cell subsets in this donor after delivery via *μVS*, indicating that our method provides advantages to viral- and mechanical-based delivery methods known to introduce expression biases in the same T cell populations.

**Figure 5.**
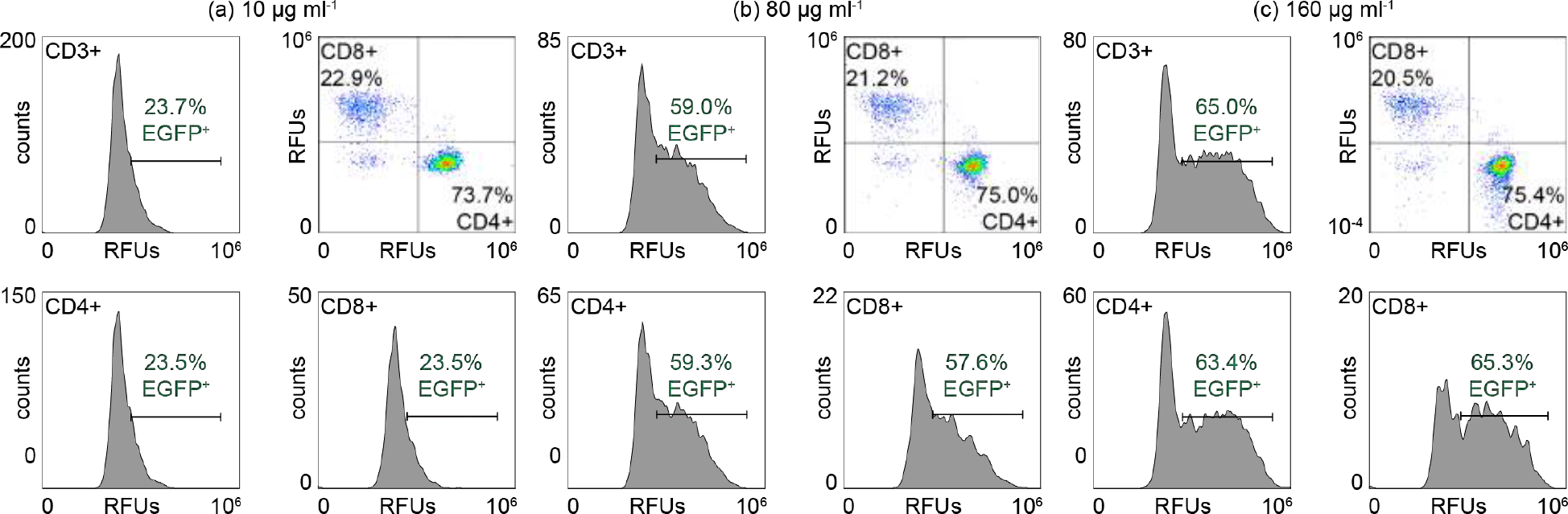
EGFP expression is equally distributed between CD4+ and CD8+ T lymphocytes after delivery of EGFP mRNA via *μVS*. Human T cells were transfected with (**a**) 10 μg mL^−1^, (**b**) 80 μg mL^−1^, and (**c**) 160 μg mL^−1^ EGFP mRNA via *μVS* and analyzed by flow cytometry for CD3, CD4, CD8, and EGFP protein expression 19 hours post transfection. Scatter plots show the percentage of CD4+ and CD8+ T cells as a percentage of the total CD3+ population. Histograms show the level of EGFP expression in CD3+, CD4+ and CD8+ T cell populations. Data represent the mean of triplicate values for each condition.

### *μVS* results in negligible perturbation of the T cell state

Electroporation and viral-based transduction of human T cells has been shown to induce significant and dramatic changes in gene expression profiles, including genes related to T cell activation and survival^11,13,39^. Therefore, we evaluated if processing cells via *μVS* would alter the expression levels of canonical T cell activation markers compared to non-processed cells. Flow analysis of device-processed and non-processed cells revealed comparable expression levels of six unique markers of T cell activation, including CD69 and CD25, 24 hours post transfection (Figure 6). Isotype controls for mouse IgG1 kappa were previously determined to have no off target or non-specific binding interactions with the T cells used in this experiment (data not shown). Therefore, these results indicate that *μVS*-based transfection of mRNA does not significantly alter the activation state of T cells 24 hours after processing.

**Figure 6.**
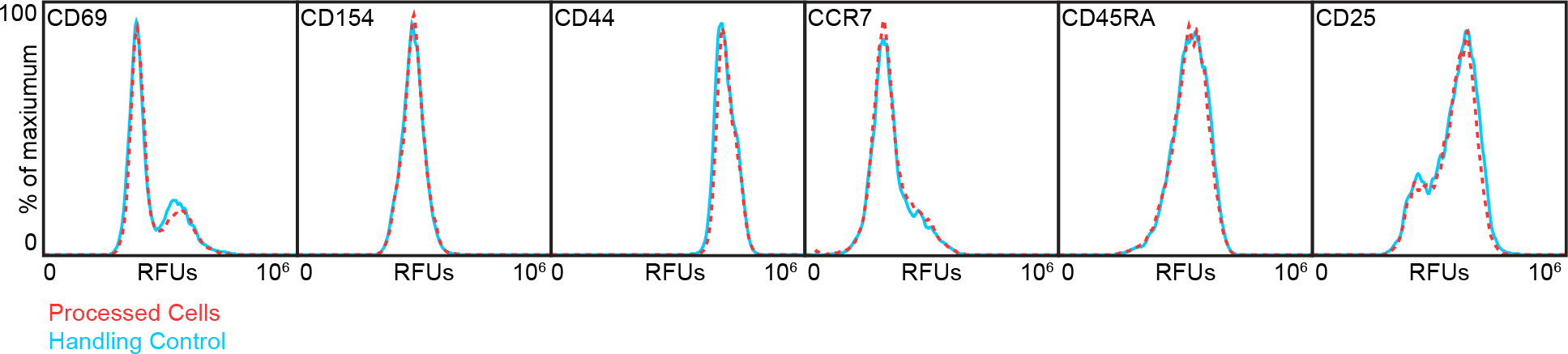
T cell activation marker expression is unaffected in cells processed via *μVS*. Flow cytometry was used to quantify the surface level expression of key T cell activation markers (CD69, CD154, CD44, CCR7, CD45RA and CD25) in T cells processed with *μVS* (processed cells, red dashed) compared to non-processed cells (handling control, blue solid) 24 hours post transfection. Histogram overlays represent expression levels as a percent of maximum value of each marker for a representative sample for each condition. Experiment was performed in triplicate.

## Discussion

The clinical and commercial standards for GMCT manufacturing require the use of viruses which is time consuming, labor intensive, expensive, and requires extensive and highly variable lead times. In addition, virally-synthesized GMCTs require significant safety, regulatory and release requirements, and strict monitoring of patients for up to 15 years after treatment^9^. Non-viral based delivery is an attractive alternative to viral GMCT manufacturing as it permits the intracellular delivery of a variety of nucleic acid constructs (DNA, RNA) and macromolecules (protein, protein complexes) into cells for permanent, persistent, or transient modification. Further, these methods provide additional advantages over the current standard including reduced costs, ease to scale, and substantially shorter lead times. Physical transfection methods, such as electroporation, are quick, simple, and efficient, resulting in greater than 90% expression of delivered constructs in human T cells and are actively being pursed for GMCT manufacturing. However, low cell viability and recovery rates associated with these methods make them less than ideal for clinical-scale production where a large number of viable, gene-modified cells is required.

*μVS* is a promising solution to these problems that may greatly reduce the time, safety concerns, and release testing requirements associated with virally-produced GMTCs. Our *μVS*-based platform can process 5.0 x 10^7^ cells mL^−1^ at high flow rates with minimal clogging. Total cell recovery after *μVS* is exceptionally high when compared to typical recovery rates of 20% after electroporation, offering a nearly 5-fold improvement. Further, *μVS* does not affect cell viability or growth rates for at least 1 week after transfection relative to non-processed cell controls. These values were significantly greater than reports of <40% viable cells and significant cell death rates 2 to 3 days after transfection by electroporation^11^. Based on a 43% yield achieved in our study, we predict that delivery of a CAR-encoding construct with similar CAR expression efficiency in 1mL of 5.0 × 10^7^ T cells would produce approximately 2.1 × 10^7^ CAR+ T cells in less than 1 minute via *μVS*. Upon activation and expansion of these cells, 3.4 × 10^8^ CAR+ T cells would be produced after four doubling times (~7-8 days post activation, doubling every other day) exceeding the total number of cells required for a single infusion of CAR-T therapy^28,29^. While future experiments to produce CAR T cells via *μVS* are ongoing, these calculations and results from our current study highlight the potential to achieve GMCT products via *μVS* by substantially reducing *ex vivo* culture times and preparing cells for infusion within 1 week of collection^40^.

As mentioned previously, electroporation is actively being explored as a potential alternative to viral-based transduction for GMCT manufacturing. However, in addition to inducing significant cell death and dramatic decreases in cell viability, electroporation can also augment the activation states and gene expression profiles of processed cells that can directly affect their survival and function *in vivo*^13^. Recently, electroporation of human T cells for Cas9-RNP-based PD-1 knockdown resulted in significant increases in cytokine secretion and dysregulation of genes related to cell proliferation and survival as a result of transfection^13^. Unlike electroporation, our examination of *μVS* revealed no change in expression of surface markers critical for the activation, function, and survival of T cells. These markers included CCR7, a chemokine receptor involved in T cell homing following antigen stimulation, CD69, an early activation marker necessary for Th17 differentiation, and the leukocyte adhesion molecule, CD44. Interestingly, changes to these markers, as well as genes involved in DNA damage, cell proliferation, and growth factor production, were detected in genetically-modified T cells as a result of electroporation and dramatically reduced their capacity to suppress tumor growth *in vivo*^11,13^. Collectively, our results indicate the *μVS*-based delivery does not perturb the native state of transfected cells, highlighting its potential to maintain the functionality and desired therapeutic effects of engineered cells upon administration.

Equal delivery and expression of desired proteins among different cell populations may be a unique feature of *μVS* and is advantageous to reduce manufacturing times for GMCTs due to the more predictable modification of subtypes (reference not needed as this is our conclusion). Lentiviral transduction efficiencies can differ among T cell subsets while electroporation is shown to adversely affect CD8+ T cells, resulting in < 40% cell viability^11^. Our study shows comparable levels of EGFP protein expression among CD4+ and CD8+ T cell subsets after delivery of mRNA via *μVS*, highlighting an additional advantage over electroporation and lentiviral delivery methods, especially for GCMT where even expression across unique T cell populations may be necessary. Further investigation of the expression profiles among T cells and other immune cells (B lymphocytes, Nature Killer cells, dendritic cells) after *μVS*-based delivery of alternative constructs, including CAR-encoding mRNA, DNA plasmids and Cas9 proteins, is currently ongoing.

Based on published reports, we observed reduced transfection efficiency with *μVS* compared to constriction-based transfection, likely due to the unique mechanical poration mechanisms facilitating membrane disruption^3^. However, this reduction is offset by exceptionally high processing rates and cell recovery obtained by the increased flow rate, small sample volume, and high cell concentration tolerances of microfluidics, while eliminating clogging issues commonly observed with microscale cell constriction^18^. Additionally, the brief time needed for *μVS* poration likely results in gentler transfection that limits cell stress and perturbation. Optimizations to the test rig, device, media composition, and constructs to further improve delivery efficiencies via *μVS* are currently ongoing. Altogether, these characteristics position *μVS* transfection as a unique microscale method suitable for generation of GMCTs at clinically-relevant scales.

As part of our product development pipeline, tube adaptors for processing volumes >10mL are currently in development to accommodate large-scale GMCT manufacturing. By processing 10 mL of cells at the above concentration, *μVS* transfection would generate the maximum therapeutic dose of CAR+ T cells (2.0 × 10^8^ cells) for Yescarta and Kymriah in less than 10 minutes^28,29^. The cost of mRNA required for a 10 mL processing volume may be prohibitive for certain autologous GMCTs, however *μVS*-based delivery could allow for immediate turnaround, making the time from apheresis to treatment less than one day if *μVS* were to be engineered into a completely closed system. We have observed that cell type and age, time post activation, and medium composition can influence debris accumulation (data not shown). Therefore, in the future we plan to optimize these parameters as we expand to larger processing volumes while increasing cell concentrations to >1.0 x 10^8^ cells mL^−1^ for synthesis of autologous and allogeneic GMCT products. We predict that *μVS*-based transfection will greatly reduce the manufacture times for clinical- and commercial-scale GMCTs, positioning this technology as a safe, sustainable and cost-effective option for GMCT production.

Taken together, our work describes a novel transfection method utilizing microfluidic post arrays and hydrodynamic conditions to facilitate the intracellular delivery of mRNA to human T cells with high transfection efficiency, cell viability, and cell recovery, and negligible perturbation to the T cell state. Our chip design contains spacing between posts approximately two-times greater than the median cell diameter leading to transfection of cells in a manner that is accommodating to cell size variability and distributions. The hydrodynamic conditions generated by processing cells in our device permit delivery of an approximately 1.0kB EGFP mRNA construct to primary human CD3+ T cells yielding 43.4 ± 2.5% recovered, viable, and EGFP-expressing T cells using 160 μg mL^−1^ mRNA at a processing speed of 2.0 × 10^6^ cells s^−1^. We found that transfection by *μVS* resulted in equal levels of EGFP expression among CD4+ and CD8+ T cells highlighting its capacity to evenly transfect unique T cell subsets. We also observed no change in T cell activation states or proliferation rates post processing via *μVS*. This is important since many of the commonly used delivery methods, including electroporation, can lead to T cell activation, exhaustion, and/or cell death^11^. Altogether these results indicate that *μVS* is an excellent alternative to current transfection methods with promising attributes for discovery-stage research, clinical development, and commercial manufacturing of engineered gene-modified human T cells.

Future work is focused on demonstrating the utility of *μVS* for research, clinical and commercial applications with an initial emphasis on CAR-T cell production. This includes evaluating delivery of different length DNA and RNA constructs, protein complexes, and co-delivery of these molecules, for applications requiring transient or stable protein expression. Supplementing flow cytometry profiling, we are also evaluating the influence of *μVS* on T cell activation and survival, function, and efficacy as a therapeutic by cytokine secretion analysis, whole transcriptome sequencing, and *in vitro* cellular assays. Ultimately, our aim is to validate *μVS*-modified cells *in vivo* to translate into scalable, cost-effective GMCT products for patients in need.

## Materials and Methods

### Device Design & Fabrication

Devices were designed with a 4.8 mm × 9.8 mm footprint and contained a 960 μm wide by a 40 μm deep flow cell. This flow cell was designed to contain a post array consisting of 40 μm diameter posts, with a pitch, or distance, between the midpoint of two adjacent posts in a post row of 60 μm orthogonal to the bulk flow direction, and a 500 μm pitch in the bulk flow direction (Figure 1c-d).

Device fabrication was achieved using industry standard semiconductor processes (Australian National Fabrication Facility, South Australia Node) and fused silica wafers. The flow cell and array geometries were constructed through anisotropic deep reactive ion etching (Figure 1c). Deep reactive ion etched flow cells were thermally bonded to a fused silica lid containing 700 μm diameter laser machined through holes for the inlet and outlet. After fabrication, device and feature geometries were verified using scanning electron microscopy (Figure 1c), white-light interferometry (not shown) and digital microscopy (not shown). Specific details on device design and fabrication are outlined in the *Supplemental Information (SI).*

### Experimental Rig Development

A purpose-built experimental rig was developed to operate a microfluidic chip (Figure 1b) using an operating pressure between 0 and 150 pounds per square inch (PSI) and measure flow rates ranging from 1 mL min^−1^ to 1 mL s^−1^. To accomplish this, compressed nitrogen was regulated down to less than 150 PSI using a calibrated two-stage regulator and filtered with a 5 μm compressed air filter (McMaster Carr, 4414K71). Compressed nitrogen flow was then controlled with a manual on/off valve (McMaster Carr, 4379K61) and volumetric flow rates were measured with a calibrated mass flow meter (Alicat Scientific, M-1SLPM-D). Compressed nitrogen was then used to pneumatically pump samples of suspended cells and constructs through the microfluidic chip. The samples were housed in a 1.5 mL tube and placed in a tube adaptor (Elveflow, KRXS) which was coupled to an in-house fixture with outlet tubing for sample collection (Figure 1d). More details regarding rig and device development, fabrication, and production are available in the *SI*.

### Hydrodynamic Characterization and Computational Fluid Dynamic Simulations

Non-dimensional equations were used to calculate the Reynolds number (*Re*) for flow in channels and around a cylindrical post^33^. This was performed to assess if flow past posts within the post array region of chips was operating within hydrodynamic condition required for vortex shedding (*Re*_*o*_ > 40) while also determining if these conditions could be the result of flow through channels. The equations are based on the device’s volumetric flow rate (Q), kinematic viscosity of the fluid (*v*) and specific device geometries.

Using the non-dimensional analysis, we calculated the Reynolds number around an object (*Re*_*o*_) where the object is a cylindrical post with a 40 μm diameter representing posts in our chip design. This allowed us to quickly and reasonably assess the flow conditions around posts.

We used two-dimensional computational fluid dynamics techniques and ANSYS Fluent to simulate hydrodynamic conditions in the unit array geometry using the free stream velocity (*v*_∞_) or the average fluid velocity within the flow cell. Simulation were performed to assess *μVS* flow development time at representative hydrodynamic conditions. *μVS* flow development time was simulated by examining the transient drag and lift coefficients acting on the posts while also visualizing velocity contours, velocity vectors and vortex identification using Q-criterion.

### Enrichment and activation of human CD_3+_ T cells

Primary human CD3+ T cells were negatively selected from single donor PBMCs using standard techniques. For revival and culture, 5.0 × 10^6^ cryopreserved T cells were expanded using CD3/CD28 T cell activator (StemCell Technologies) and standard conditions for 16 days to achieve sufficient cell numbers for the experimental workflow. Further details outlined in *SI*.

### EGFP mRNA delivery into primary T cells via *μVS*

All solutions processed though the device and chip were filtered prior to use using a 0.22 μm filter to remove particulates. For on chip processing, T cells were removed from culture, washed, resuspended in processing medium consisting of Immunocult-XF media (StemCell Technologies), 25 mM trehalose dihydrate NF (JT Baker, VWR) and 5% v/v DMSO (Corning Cellgro, Fisher Scientific), and filtered using a sterile 40 μm filter. Each mRNA condition was evaluated in triplicate at a final cell concentration of 1.6 × 10^7^ cells mL^−1^ and 400 uL processing volume.

The sample rig and tubing were sterilized before use via 70% ethanol wipe down and flush. Immediately before processing, each sample was mixed in a 1.5mL tube with the appropriate volume of EGFP mRNA (996 nucleotides translated into a 26.9 kDa protein, 1 mg mL^−1^ L-7601, TriLink BioTechnologies) at final concentrations ranging from 10 μg mL^−1^ to 160 μg mL^−1^ (30 nM to 473 nM). The sample was mixed thoroughly, mounted in the tube fitting, and exposed to 120 PSI nitrogen pressure to drive the sample through the chip and induce intracellular mRNA uptake via *μVS* (Figure 1a). Processed samples were then collected in a 15 mL conical tube and placed on ice until the completion of the experiment with a maximum time on ice of less than 4 hrs. After each run, the rig and tubing were flushed with 70% ethanol and a new microfluidic chip was placed in the rig. Time equals zero for mRNA expression started when all samples were processed, removed from ice, and returned to culture medium. Control samples were set up in triplicate and remained at room temperature while the experimental samples were processed before being placed on ice. Control samples that were not device processed consisted of 1.6 × 10^7^ cells mL^−1^ in processing medium (handling control) and processing medium containing 160 μg mL^−1^ mRNA (mRNA control). Controls were used to normalize the cell viability and recovery for the experimental samples. Additional device control samples were set up at 1.6 × 10^7^ cells mL^−1^ in processing medium and processed through the device to determine the impact of *μVS* on cell survival and state without additional external factors.

### Post processing cell culture analysis

After the last sample was processed and placed on ice, samples were removed for post processing cell viability and counts. The remaining samples were diluted in X-VIVO10 (04-380Q, Lonza) at a concentration of approximately 8.0 × 10^5^ cells mL^−1^ with 100 IU mL^−1^ IL-2 (200-02, Peprotech) and cultured in 6 well non-TC treated plates (Corning) at 37 °C in 5% CO_2_ for growth, viability, and activation marker and EGFP expression analysis. Additional IL-2 at 100 IU mL^−1^ was added on days 2 and 4 after transfection and cells were discarded on day 7.

Initial T cell viabilities and post processing concentrations were used to determine the recoveries and yields. Cell growth and viability of each sample in X-VIVO10 with 100 IU mL^−1^ IL-2 was monitored over a period of seven days using a Countess II automated cell counter with trypan blue dye exclusion. EGFP expression and persistence at various time points post transfection was monitored using flow cytometry (Attune Nxt, ThermoFisher) with 1 μM propidium iodide (R37169, Sigma Aldrich) to exclude dead cells.

Expression efficiency along with cell recovery and cell viability were enumerated as a function of mRNA concentration to determine the mRNA concentration that results in the ‘yield’ of recovered, viable and transfected cells defined as: *y* = *rve* where *y* is yield, or the fraction of recovered, viable, and transfected cells or percent of input cells that remained viable and expressed EGFP after device transfection. *r* is the fraction of recovered cells after device processing, *v* is the viability of the recovered cells, and *e is* the efficiency of transfection or the fraction of viable cells expressing EGFP. Yield and efficiency of EGFP expression were calculated using the highest EGFP expression value from cultures, which occurred at approximately 19 hrs post processing.

To assess the impact of *μVS*-based mRNA delivery on T cell activation, one sample from each replicate in the control and device processed (no mRNA) groups were labeled with fluorescent labeled antibodies against various activation markers 24 h post transfection. Samples were rinsed and re-suspended in flow buffer containing 25 μL mL^−1^ of each of the following monoclonal anti-human antibodies per sample: anti-human CD3-AF700 (56-0037-42, ThermoFisher), - CD40L/CD154-FITC (11-1548-42, ThermoFisher), -CD25-PE (120257-42, ThermoFisher), -CCR7-APC-eF780 (47-1979-42, ThermoFisher) -CD44-APC-eF780 (47-0441-80, ThermoFisher), -CD69-eF450 (48-0699-42, ThermoFisher), and -CD45RA-SB702 (67-0458-42, ThermoFisher). The samples were incubated on ice for 30 minutes, rinsed, and analyzed via flow cytometry.

### EGFP expression levels in CD_3+_ T cell subsets

Expression efficiency was examined in CD4+ and CD8+ T cell subsets using fluorescently-labeled monoclonal antibodies and flow cytometry analysis 27 h post transfection. Samples were removed from each of the 10, 80, and 160 μg mL^−1^ EGFP mRNA cultures, rinsed and re-suspended in 100 μL of 25 μg mL^−1^ mouse anti-human CD3-AF700 (56-0037-42, ThermoFisher), CD4-PE-Cy7 (25-0048-42, ThermoFisher), and CD8-SB600 (63-0088-42, ThermoFisher) in DPBS containing 1% bovine serum albumin and 2 mM EDTA (flow buffer) to quantify the percentage of CD4+ and CD8+ T cells expressing EGFP. Labeled cells were analyzed via flow cytometry and compensated using AbC compensation beads (A10497, Invitrogen) labeled with the above antibodies and EGFP-expression cells.

## Data Availability

The datasets generated and/or analyzed in the current study are available from the corresponding authors by reasonable request.

## Acknowledgements

This work was funded in part by IndieBio (indiebio.co), SOSV (sosv.com), Jobs for NSW (jobsfornsw.com.au), Y Combinator (ycombinator.com), AusIndustry, Social+Capital, Main Sequence Venture, Founders Fund, MTP Connect, NSW Health Medical Device Fund and angel investors. The authors would also like to acknowledge Simon Doe, Hamish Hawthorn, Ben Wright, Mike Nicholls, Warren McKenzie, Todd Martin, Niranjana Nagarajan, Steve Gourlay, Mike Bowles, Geoff Facer, and Heidi Hagen for their guidance, advice and support to pursuing science in the startup environment. Single donor human primary T cells were provided as a gift by Eureka Therapeutics, Inc. The content is solely the responsibility of the authors and does not necessarily represent the views of anyone acknowledged in this section.

## Author contributions

Conception, experimental design, simulations, analysis and interpretation were performed by J.A.J, A.A.T, K.H.W.J.L, M.N.K., A.A.L. and R.S.P. The manuscript was drafted by J.A.J, A.A.T, K.H.W.J.L., M.N.K, R.S.P. Revisions to the manuscript were provided by J.A.J., M.N.K., A.A.L., J.A., C.P., J.N., D.G. and R.S.P.

## Supplemental Methods

### Device Design & Fabrication

The microfluidic chips were produced with deep reactive ion etching (DRIE) and thermal bonding. Vertical sidewalls were a product of DRIE on fused silica and a bulk material bond was produced with a thermal bond (Figure 1b). This production method allowed for high operating pressures and the use of an optically-transparent fused silica substrate allowed for device inspection.

**SI Table 1.** Summary of designed and fabricated device features

| Feature | Design | Fabrication | Difference |
| --- | --- | --- | --- |
| Flow cell height ($h_{fc}$) | 40.0 $\mu\text{m}$ | 40.9 $\mu\text{m}$ | 2.3% |
| Post diameter ($d$) | 40.0 $\mu\text{m}$ | 42.2 $\mu\text{m}$ | 4.0% |
| Row pitch ($p_r$) | 60.0 $\mu\text{m}$ | 59.5 $\mu\text{m}$ | -0.8% |
| Channel wall thickness ($t$) | 6.0 $\mu\text{m}$ | 6.2 $\mu\text{m}$ | 3.3% |

The use of semiconductor processes resulted in minimal variation between designed and fabricated device geometries. The device yield, or percentage of successfully fabricated devices, was typically 95%. This high yield is attributed to relatively simple device geometries. For example, the maximum fabricated feature aspect ratio was approximately 6.6. DRIE is frequently used to generate features with substantially higher aspect ratios^1^.

The devices were designed with inlet and outlet channels that created uniform flow conditions^2^ and improved cell recovery. In preliminary experiments, device designs with open or channel-free inlets and outlets were capable of intracellular delivery; however, cell recovery was significantly lower and cell debris or buildup on the upstream side of the post was observed. These preliminary device designs were also prone to clogging when used to process larger volumes of cells and at high cell densities. This clogging was attributed to mechanical lysis of cells after impacting a post.

The introduction of inlet and outlet channels substantially reduced the opportunity for this form of cell lysis along with related device clogging. For the specific design used herein, fabricated posts occupied approximately 68.2% of the flow cell width, whereas the fabricated inlet and outlet channel structures occupied approximately 10.3% of the same flow cell width. Cellular debris was still observed at the inlet of the channels after processing, but cell recovery rates remained high and the devices were less prone to clogging particularly at the cell densities evaluated in this study.

### Experimental Rig Development

Microfluidic equipment is typically limited to low flow rates, small sample volumes, and low operating pressures. Thus, a custom test rig was developed to withstand the high pressures required to generate *μVS* within the microfluidic chip. Direct measurement of microfluidic flow rates were not feasible with readily available commercial equipment due to the (1) significant flow rates, (2) sensitive nature of primary T cells, and (3) short flow times. For example, a 400 μL sample containing 16 × 10^6^ cells mL^−1^, processed at an operating pressure of 120 psig (8.2 ATM) resulted in a measured flow rate between 5 to 8 mL min^−1^ and total sample flow time of approximately 3 seconds. Additionally, the brief sample flow time coupled with limited tubing lengths, between the mass flow meter and the chip fitting, meant artifacts caused by diffusion and permeation of compressed nitrogen through the tubing wall was negligible, relative to flow of nitrogen driving the sample through the tubing and chip. Direct microscopy imaging of *μVS* with the current experimental rig was not possible because of the flow rate, short cell residence time, and rapid transfection.

This purpose-built system allowed us to quantify flow rates without disrupting (1) the hydrodynamic conditions within the chip or (2) the sample flow through the tubing. During preliminary experiments, it was observed that significant bends and kinks in the tubing would adversely affect cell viability and recovery – this was attributed to high shear recirculation regions created within the deformed tubing. Careful selection of tube inner diameter, length and material were required to generate the ideal flow conditions. Small changes in the inner diameter of the tubing resulted in significant changes in volumetric flow rates and hydrodynamics conditions within the chip, while excessive tubing lengths increased total sample loss. Cumulatively, an experimental rig was engineered to provide quantitative flow measurements of flow paths and subject cells to precisely-controlled hydrodynamic conditions within the microfluidic chip.

### Hydrodynamic Characterization

Non-dimensional equations were used to calculate the Reynolds number (*Re*) in channels and for flow around a cylindrical post^2^. This was done to determine if the post arrays were operating in the hydrodynamic conditions that support vortex shedding (*Re*_*o*_ > *40*). The equations are based on the device’s volumetric flow rate (*Q*), kinematic viscosity of the fluid (ν) and specific device geometries.

Using non-dimensional analysis, the following Reynolds numbers were calculated: flow cell (*Re*_*fc*_), inlet channels (*Re*_*c*_), gap between posts (*Re*_*g*_), and flow around an object where the object is a cylindrical post (*Re_*o*_*). This allowed us to quickly and reasonably assess (1) if flow conditions were hydrodynamic and (2) the most probable cause of hydrodynamic conditions. *Re*_*fc*_, *Re*_*c*_, and *Re*_*g*_ were calculated as follows: 

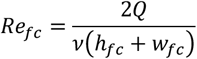

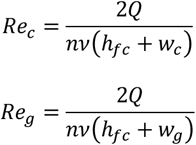

Where *Q* is the device volumetric flow rate, *h*_*fc*_ is the height of the flow cell, and *v* is the kinematic viscosity of the fluid; the respective widths of the flow cell, channel, and gap are *w*_*fc*_, *w*_*c*_, and *w*_*g*_; and *n* is the number of gaps or channels and is calculated by: 

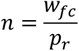

Where *p*_*r*_ is the row pitch. *Re*_*o*_ was calculated as follows:

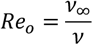

where *v*_∞_ is the free stream velocity and *d* is the post diameter. *v*_∞_ was calculated as follows: 

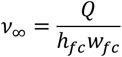

*Re*_*o*_ was then used to calculate the Strouhal number (*St*) for a smooth cylinder^1^ using the following equations: 

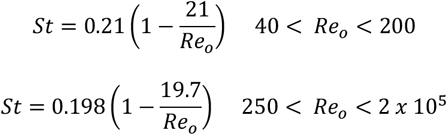

*St* was then used to approximate the frequency (*f*) of vortex shedding using the following equation: 

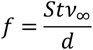

### Pan T cell Culture

Cryopreserved primary human CD3+ T cells were negatively selected from single donor primary blood mononuclear cells (PBMCs) using standard bead-based techniques. For revival and culture, 5 × 10^6^ pan T cells cell were thawed and seeded in X-VIVO10 with gentamycin and phenol red (Lonza). The cells were activated using CD3/28 T cell activator solution (StemCell Technologies) and 100 IU mL^−1^ recombinant human IL-2 (PeproTech) on the day of thaw and cultured at 37 °C in 5% CO_2_. Pan T cells were expanded for 16 days with addition of medium and IL-2 (100 IU mL^−1^) every two to three days to maintain cell concentration at or below 1 x 10^6^ cells mL^−1^. All data was collected between days 17 and 24 post thaw and activation.

### EGFP mRNA delivery to primary T cells at different concentrations

All solutions that were used on chip were filtered using 0.22 μm filtration prior to use to remove particulates that could lead to clogging. For on chip processing, T cells were removed from culture and pelleted via centrifugation (5 min at 300 xg). Supernatant was removed via aspiration and the cell pellet was resuspended in 1× Dulbecco’s phosphate buffered saline (DPBS, Gibco). Cell were subsequently pelleted via centrifugation and aspirated. Rinsed cells were resuspended in filtered processing medium composed of Immunocult-XF (StemCell Technologies), 25 mM trehalose dihydrate NF (JT Baker, VWR) and 5% v/v DMSO (Corning Cellgro, Fisher Scientific) at a cell concentration around x 10^7^ cells mL^−1^. Cell concentration was enumerated using a Countess II automated cell counter (Invitrogen) and trypan blue dye exclusion to assess viability. Three replicate samples were evaluated for each condition. To make the samples, 6.4 x 10^6^ cells were removed from the stock cell suspension, placed in 1.5 mL tubes, and diluted with processing medium to a final volume of 400 μL minus the mRNA volume. mRNA was added immediately before processing for 1.6 × 10^7^ cells mL^−1^ final cell concentration. Samples were filtered in two separate batches (160 μg mL^−1^ mRNA control, 10 μg mL^−1^, 40 μg mL^−1^, and 80 μg mL^−1^ as the first set of samples, handling control, 120 μg mL^−1^, 160 μg mL^−1^ and no mRNA device processed control as the second set) to reduce cell clumping, cluster formation, and potential for chip clogging.

The sample rig and tubing were sterilized before use with 70% ethanol wipe down and flush between each run. Prior to processing, each sample was mixed with the appropriate volume of EGFP mRNA (996 nt translated into a 26.9 kDa protein, 1 mg mL^−1^ TriLink BioTechnologies, San Diego CA, L-7601) at final concentrations ranging from 10 μg mL^−1^ to 160 μg mL^−1^ (30 nM to 473 nM). Sample was mixed thoroughly by pipetting, sample tube was mounted in the tube fitting, and the fitting clamped. Cell and mRNA suspension was then exposed to 120 PSIG nitrogen pressure to drive the sample through the chip and induce intracellular mRNA uptake via *μVS* (Figure 1a). Processed samples were collected in 15 mL conical tubes and placed on ice until for the duration of the experiment to sync expression time points between samples. After each run, the rig and tubing were flushed with 70% ethanol and a new microfluidic chip was placed in the rig. Time equals zero for mRNA expression started when all samples were processed, removed from ice, and returned to culture medium. All samples within a set were processed within 30 min and the total time on ice for the samples ranged from 0.5 to 3.75 hrs. Control samples were performed in triplicate and remained at room temperature while the experimental samples were processed. Control samples that were not processed through the device consisted of 1.6 × 10^7^ cells mL^−1^ in processing medium (handling control), which was used to normalize the cell viability and recovery for the experimental samples, and in processing medium containing 160 μg mL^−1^ mRNA (mRNA control). Additional device control samples were set up at 1.6 × 10^7^ cells mL^−1^ in processing medium and ran through the device to determine the impact of *μVS* on cell survival without additional external factors (processing control).

After the last sample was processed and incubated on ice for 5 min, all samples were removed from ice, re-suspended, and an aliquot was taken from each sample for post processing cell viability and concentration quantitation. The remaining cells were diluted 1:20 in X-VIVO10 at a concentration of 8 × 10^5^ cells mL^−1^ with 100 IU mL^−1^ IL-2. Cultures were added to 6 well non-TC treated plates and cultured at 37 °C in 5% CO_2_ for growth, activation marker, and EGFP expression analysis at later times. Cell viability was monitored using trypan blue dye exclusion and growth was quantified using the Countess II cell counter. Additional IL-2 was added on days 2 and 4 after transfection and the cells were discarded on day 7, at a culture age of 24 d post thaw and activation.

Initial T cell viabilities and post processing concentrations were quantified by counting a minimum of two aliquots from each sample. Counts were used to determine the recovery and yield shown in Figure 2a. Cell growth and viability of each sample in growth medium was monitored over seven days (Figure 3a-b). EGFP expression and persistence at different time points post transfection were monitored using flow cytometry (Attune NxT flow cytometer) and propidium iodide (1 μM final concentration, Sigma Aldrich) to exclude dead cells (Figure 3c).

### Quantifying EGFP expression levels in T cell subsets

Expression efficiency was examined among CD4+ and CD8+ T cell subsets using fluorescently-labeled monoclonal antibodies and flow cytometry. Cultures were analyzed 27 hr after being returned to culture medium. Aliquots were collected from 10, 80, and 160 μg mL^−1^ cultures, placed in a v bottom 96 well plate, diluted with DPBS, pelleted via centrifugation, and aspirated. Cells were resuspended in 100 μL mL^−1^ of 25 μL mL^−1^ each mouse anti-human CD3-Alexafluor700 (ThermoFisher PN 56-0037-42), mouse anti-human CD4-PE-Cy7 (ThermoFisher PN 25-0048-42), and mouse anti-human CD8a-Super Bright 600 (ThermoFisher PN 63-0088-42) in DPBS containing 1% bovine serum albumin and 2 mM EDTA (flow buffer). Antibody staining was used to quantify the percentage of CD4+ and CD8+ T cells expressing EGFP. Cells were incubated in the antibody cocktail on ice for 30 minutes, then diluted with 100 μL flow buffer, pelleted via centrifugation, and aspirated. Samples were resuspended in 200 μL flow buffer, pelleted via centrifugation, and aspirated a second time before resuspending in 200 μL flow buffer and analyzed via flow cytometry. The experiment was compensated using a combination of AbC compensation beads (ThermoFisher) labeled with each antibody and EGFP-expressing cells (BL1 channel).

### Medium composition

The impact of exposing human PBMCs to DMSO for long periods of time in culture conditions (37 °C, 5% CO_2_), was previously described by de Abreu Costa *et al.*^*3*^ Briefly, long term exposure to various concentrations of DMSO resulted in increased cell death, decreased proliferation, and reduced cytokine production with increasing concentrations of DMSO and long exposure times (>4 h), after activation with phorbol-12-myristate-13-acetate (PMA)^3^. In our study, we showed that DMSO at 5% v/v did not impact cell viability until PBMCs were incubated in this solution for greater than 24 hrs at 37 °C and 5% CO_2_. Cytokine production in T cells after activation with PMA was negatively impacted by exposure to 5% DMSO but required eight hours of incubation time to show an effect. The max time for samples in processing medium was under four hours, and the samples were held on ice. The cells were processed in the order of lowest to highest mRNA concentration meaning 10 μg mL^−1^ spent the longest time on ice exposed to DMSO post processing. This could be why the growth rate of this group of cells was reduced post processing compared to the other groups, meaning that this reduction in growth rate is an artifact due to the experiment length. The decrease in viability of all groups compared to the handling control may be due to increased cell death upon return to culture due to unsuccessful membrane repair attempts, bulk mechanical lysis along with thermal shock, and medium stress, but this viability decrease is small when considering the device processed control.

Processing medium ionic strength may also have an impact on the overall ease of pore formation and cell recovery after *μVS* exposure. We observed a trend of decreasing viability with increasing mRNA concentrations, though this decrease was slight. This was potentially due to the higher percentage of mRNA solution added to samples containing higher concentrations of mRNA. For example, a 10 μg mL^−1^ sample contained 1% v/v mRNA buffer in the processing medium, while 160 μg mL^−1^ sample contained 16% v/v mRNA buffer. The mRNA solution is at a low ionic strength buffer (1 mM sodium citrate) relative to Immunocult-XF. This indicated that the reduction in total processing medium ionic strength associated with increasing mRNA concentration is likely cause of decreasing viability. The presence of mRNA in solution also decreased the concentration of trehalose, which could also result in lower cell viability with increasing mRNA concentration. Exploration of these variables, as well as the addition of concentrated buffers to adjust ionic strength, are currently ongoing.

## Supplemental Figures

**Supplemental Figure 1.**
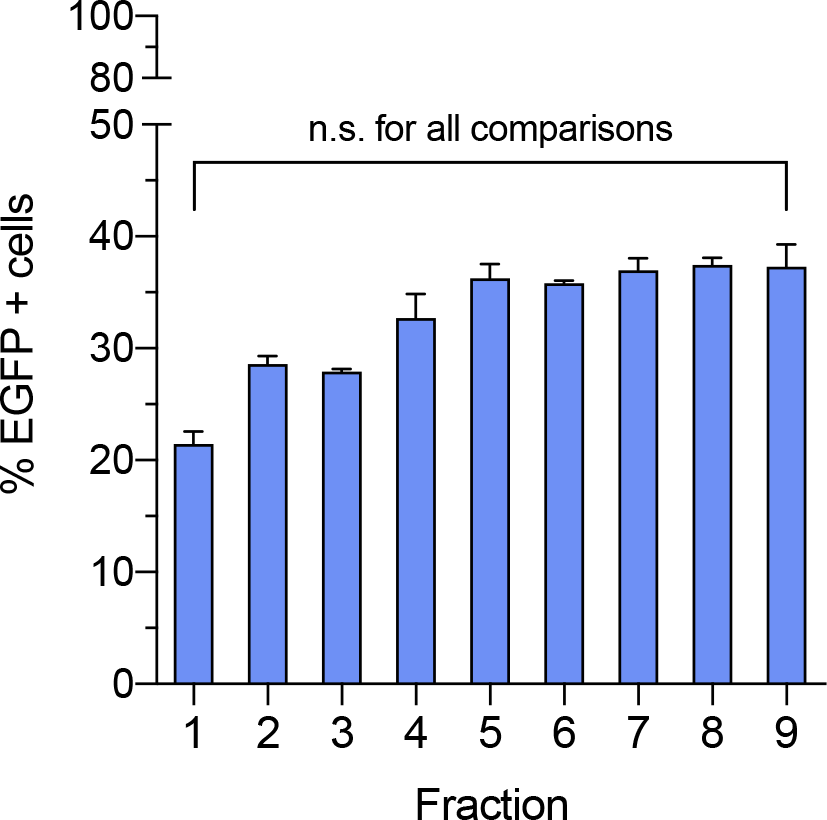
Consistent EGFP expression through duration of *μVS* transfection. Levels of EGFP protein expression were quantified in human T cells 25 hours post transfection with 80 μg mL^−1^ EGFP mRNA via *μVS*. Fractions represent post-processing cell samples collected at various time points from the start (Fraction 1) and end (Fraction 9) of the experiment. Data represent the mean ± SD of triplicate values. P values were calculated between groups by oneway ANOVA and post-hoc Kruskal-Wallis multiple comparisons test.

**Supplemental Figure 2.**
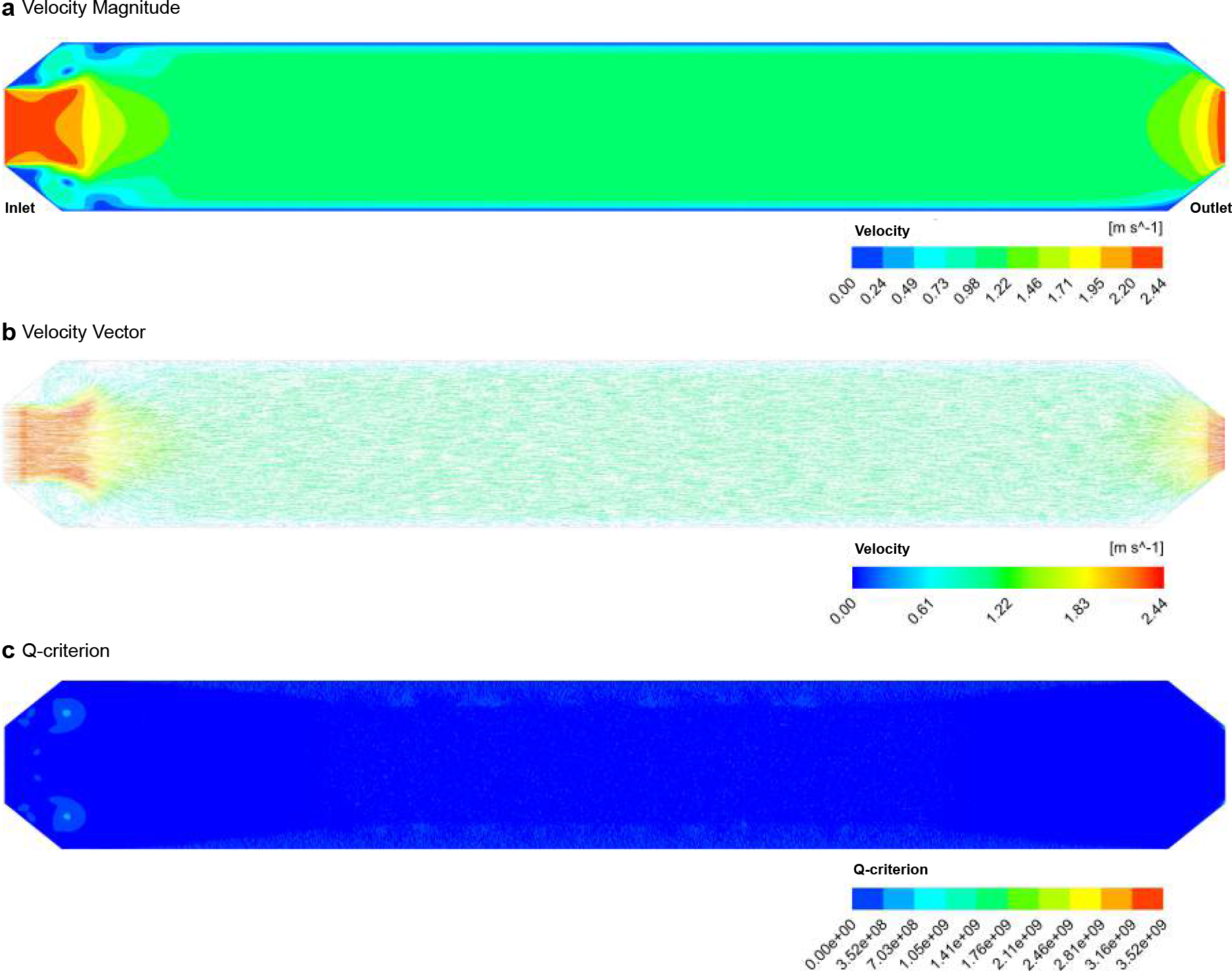
Computational fluid dynamic simulation of an empty chip. Simulation of the flow dynamics in a 960μm width × 6.95mm length chip without posts depicting (a) velocity magnitude (m s^2^), (b) velocity vector (m s^2^), and (c) Q-criterion. Flow direction was simulated from left to right.

## Supplemental Table

**Supplemental Table 1.**
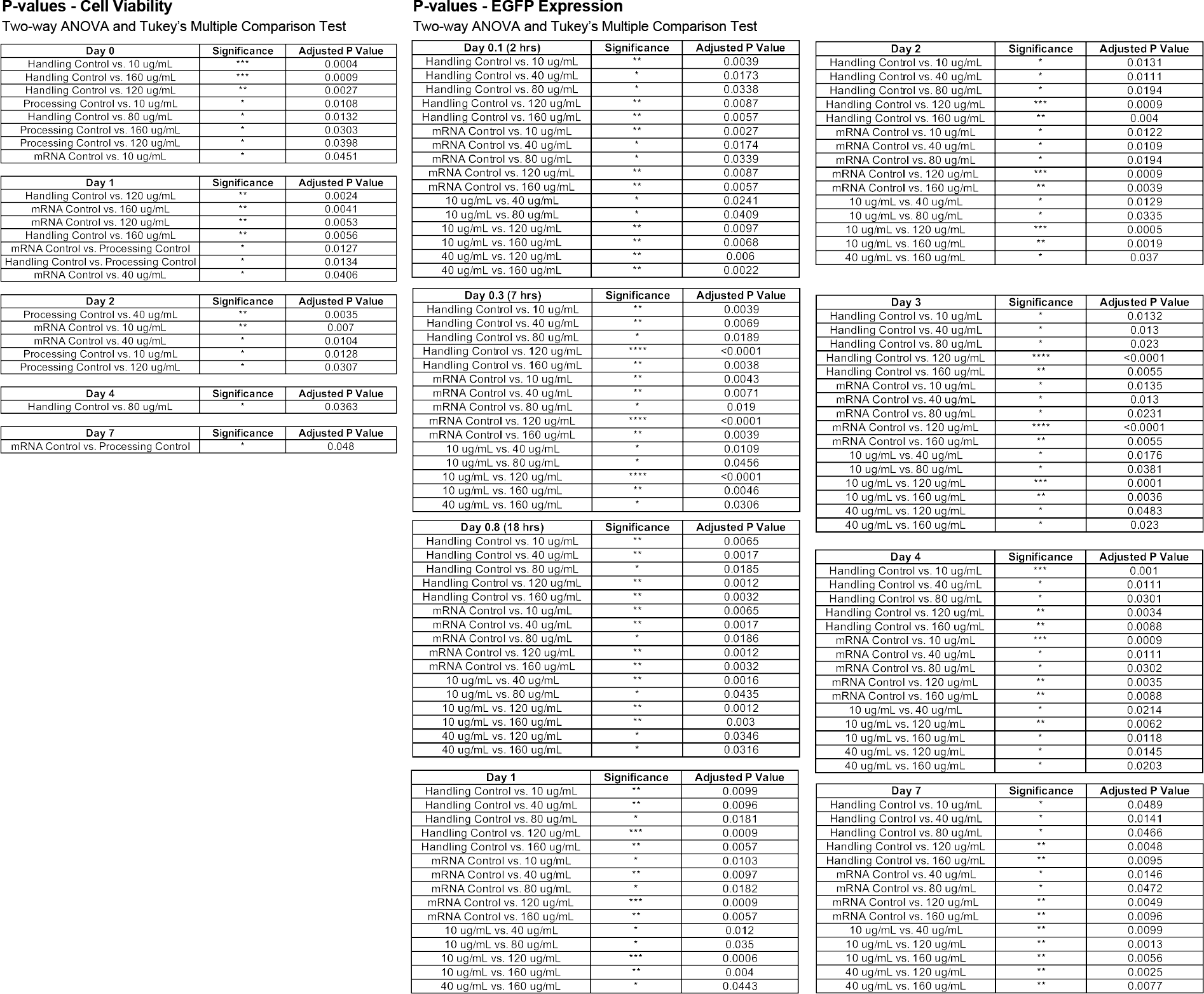
P-values for cell viability and EGFP expression efficiency measurements from Figure 4.

